# Identification of specific biomarkers and pathways in the synovial tissues of patients with osteoarthritis in comparison to rheumatoid arthritis

**DOI:** 10.1101/2020.10.22.340232

**Authors:** Hanming Gu, Gongsheng Yuan

## Abstract

Osteoarthritis and rheumatoid arthritis are two common arthritis with different pathogenesis. Here, we explore the difference of genes and biological pathways in human synovial fibroblasts by using a bioinformatics method to clarify their potential pathogenesis. The GSE7669 dataset was originally produced by using an Affymetrix Human Genome U95 platform. We used the KEGG and GO analysis to identify the functional categories and pathways. Our results suggested that biological adhesion and cell adhesion are the main signaling pathways in osteoarthritis in comparison to rheumatoid arthritis. Furthermore, Albumin, MAPK3, PTPRC, COL1A1, and CXCL12 may be key genes in osteoarthritis. Therefore, our study provides potential targets for the specific and accurate therapy of osteoarthritis.

## Introduction

Osteoarthritis (OA) is a heavy health burden for individuals, health departments, and economics^1–3^. The study of OA was mainly focused on articular cartilage, which actually had no direct association with the development of diseases^4^. Recently, the synovium and its inflammatory responses have been proved to become the targets for future therapy^5, 6^.

OA is generally caused by abnormal forces on the joints, and it is followed by the numerous production of proinflammatory cytokines, such as IL1, TNFα and IL6 in synovium^7–9^.OA and rheumatoid arthritis (RA) both can cause cartilage damage which is the most common feature between them^10^. The cartilage damage in OA is caused by the intrinsic activity of the chondrocytes^11^. The damage is regulated by the disordered production of matrix-degrading enzymes from chondrocytes^12^. In comparison, RA can cause systemic and local autoimmune reactions which lead to the activation of synovial cells to attack the cartilage^13–15^. Other common mechanisms have also been shown to operate in OA and RA. For instance, the enzymes that regulate the degradation of cartilage show numerous overlaps between them, such as MMPs^16, 17^. Moreover, the profiles of inflammatory mediators overlap between RA and OA in different conditions^5^. Recently, the development of therapy for OA is mainly dependent on the specific protein structure. OA drug production is widely dependent on genomics^18, 19^. Therefore, to identify specific targets for OA patients by genomics is of great importance.

In this study, we investigated the difference between RA and OA on human synovium. We identified the differentially expressed genes (DEGs) and the potential signaling pathways by utilizing comprehensive bioinformatics analyses. The functional enrichment, pathway analysis, and protein-protein interaction were performed to find the specific OA gene nodes in comparison to RA. These key genes could be critical and specific to guide future clinical and therapeutic interventions for OA.

## Methods

### Data resources

Gene profile dataset GSE7669 was downloaded from the GEO database (http://www.ncbi.nlm.nih.gov/geo/). The data was produced by performing an Affymetrix Human Genome U95 Version 2 Array (BIOTEC, Tatzberg. Dresden, Germany). The GSE7669 dataset contained data including 6 patients with OA and 6 patients with RA.

### Data acquisition and preprocessing

The raw data between OA samples and RA samples were analyzed by R script as described previously^20, 21^. A classical t test was used to identify DEGs with P<.05 and fold change ≥1.5.

### Gene ontology (GO) and pathway enrichment analysis

Gene ontology (GO) analysis and the Kyoto Encyclopedia of Genes and Genomes (KEGG) database were used for systematic analysis of gene functions and pathways. GO and KEGG analysis of DEGs in this study were analyzed by the Database for Annotation, Visualization, and Integrated Discovery (DAVID) (http://david.ncifcrf.gov/) and Reactome Pathway Database (https://reactome.org/). P<.05 and gene counts >10 were considered statistically significant.

### Module analysis

The Molecular Complex Detection (MCODE) was used to analyze the potential PPI networks. The function and pathway enrichment analyses were performed by using Reactome Pathway Database (https://reactome.org/), and P<.01 was used as the cutoff criterion.

## Results

### Identification of DEGs of OA in comparison to RA

To gain the insights on the difference between RA and OA, the modular transcriptional signature of OA synovial tissues was compared to that of the RA. A total of 726 genes were identified to be differentially expressed in OA samples with the threshold of P<0.05. The top 10 up- and down-regulated genes for OA and RA samples are list in Table 1.

**Table 1.**
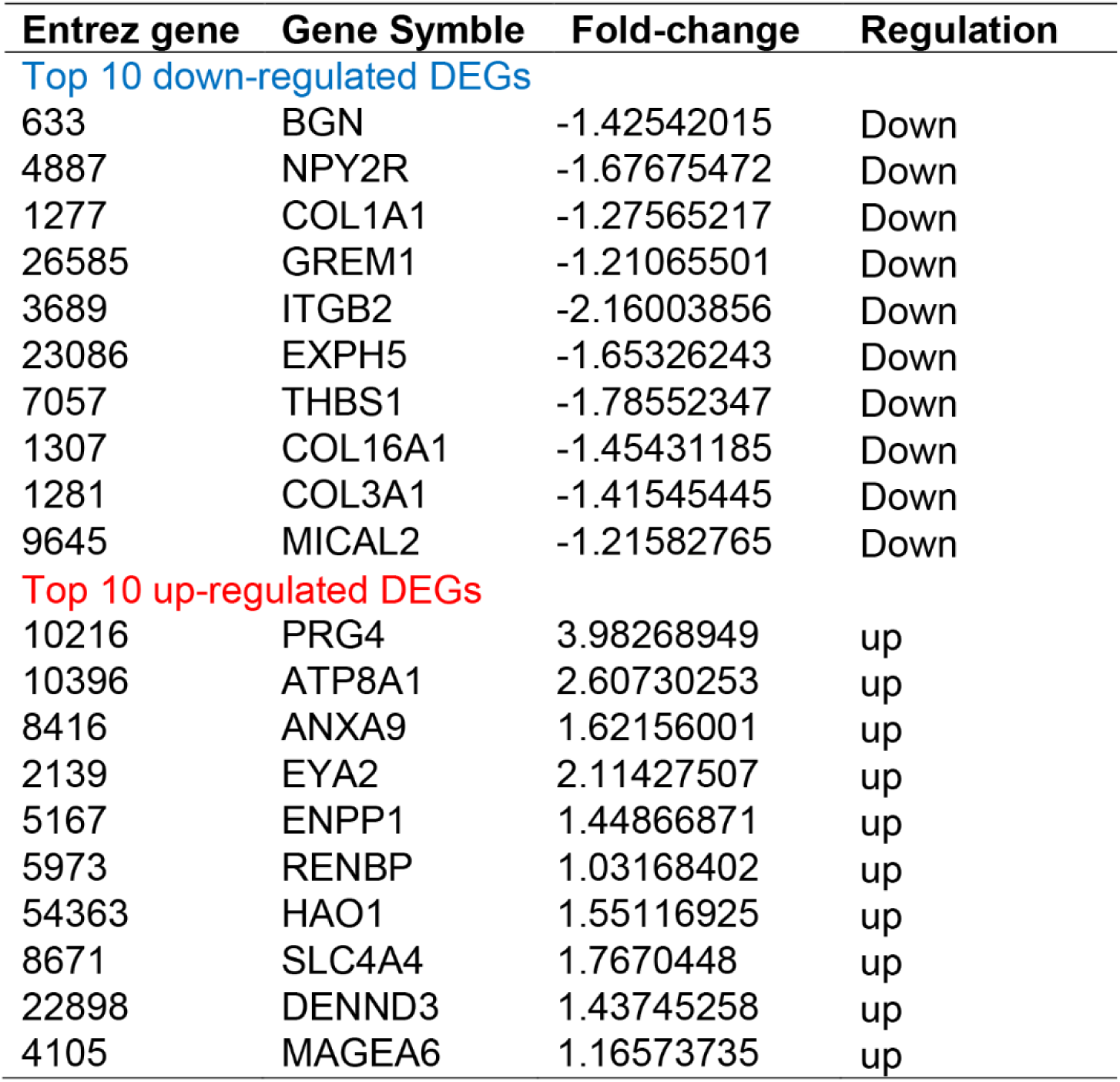

### Enrichment analysis of DEGs in OA in comparison to that in RA

To further analyze the biological roles and potential mechanisms of DEGs of OA and RA, we performed KEGG pathway and GO categories enrichment analysis. KEGG pathways (http://www.genome.jp/kegg/) is an encyclopedia for understanding genes by high-level functions. Our study showed top three enriched KEGG pathways including “Vascular smooth muscle contraction”, “Focal adhesion” and “ECM-receptor interaction” (Figure 1).

**Figure 1.**
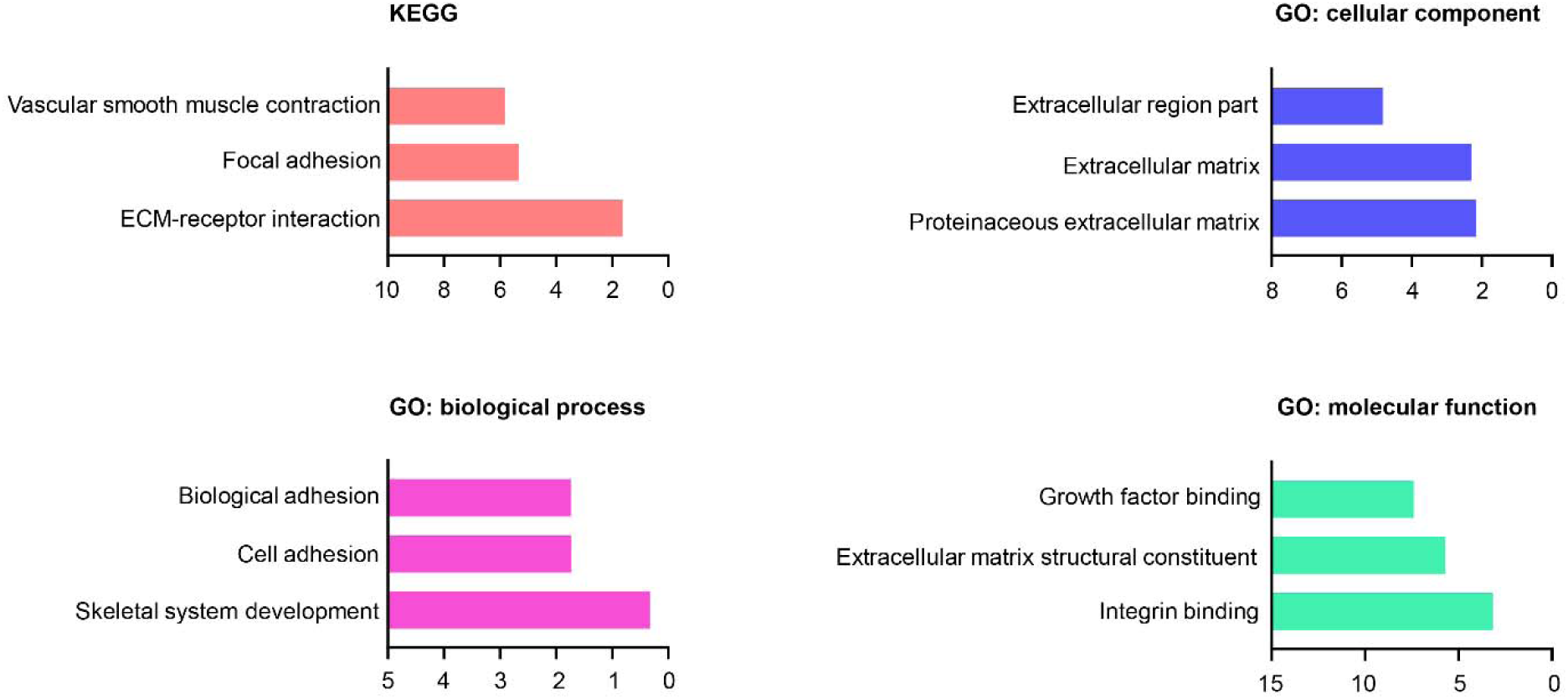
The KEGG pathways, biological process, cellular component, and molecular function terms enriched by the DEGs between OA and RA. DEGs = differentially expressed genes, KEGG = Kyoto Encyclopedia of Genes and Genomes.

Gene ontology (GO) analysis is a critical bioinformatic initiative to unify the representation of gene and gene product. We identified top three cellular components including “Extracellular region part”, “Extracellular matrix” and “Proteinaceous extracellular matrix” (Figure 1). We then identified top three biological processes: “Biological adhesion”, “Cell adhesion” and “Skeletal system development” (Figure 1). We also identified top three molecular functions: “Growth factor binding”, “Extracellular matrix structural constituent” and “Integrin binding” (Figure 1).

### PPI network and Module analysis

The PPI network was created to further explore the relationship of DGEs at the protein level. We set the criterion of combined score >0.7 and created the PPI network by using the 558 nodes and 2537 interactions between OA and RA samples. Among these nodes, the top ten genes with highest scores are shown in Table 2.

**Table 2.** Top ten genes demonstrated by connectivity degree in the PPI network

| Gene Symbol | Gene title | Degree |
| --- | --- | --- |
| ALB | albumin | 110 |
| MAPK3 | mitogen-activated protein kinase 3 | 76 |
| BMP4 | bone morphogenetic protein 4 | 58 |
| PTPRC | protein tyrosine phosphatase receptor type C | 55 |
| COL1A1 | collagen type I alpha 1 chain | 53 |
| CXCL12 | C-X-C motif chemokine ligand 12 | 44 |
| POSTN | periostin | 44 |
| COL3A1 | collagen type III alpha 1 chain | 43 |
| NCAM1 | neural cell adhesion molecule 1 | 43 |
| BGN | biglycan | 42 |

The top two significant modules of OA versus RA samples were selected to depict the specific functional annotation of genes (Figure 2 and 3). We identified top five signaling pathways in module 1: Post-translational protein phosphorylation, Regulation of Insulin-like Growth Factor (IGF) transport and uptake by Insulin-like Growth Factor Binding Proteins (IGFBPs), Extracellular matrix organization, ECM proteoglycans, and Molecules associated with elastic fibers. We also identified five top signaling pathways in module 2: Collagen chain trimerization, Collagen degradation, Assembly of collagen fibrils and other multimeric structures, Integrin cell surface interactions, Extracellular matrix organization by using Reactome Pathway Database (https://reactome.org/) (Supplemental Table S1 and S2).

**Figure 2.**
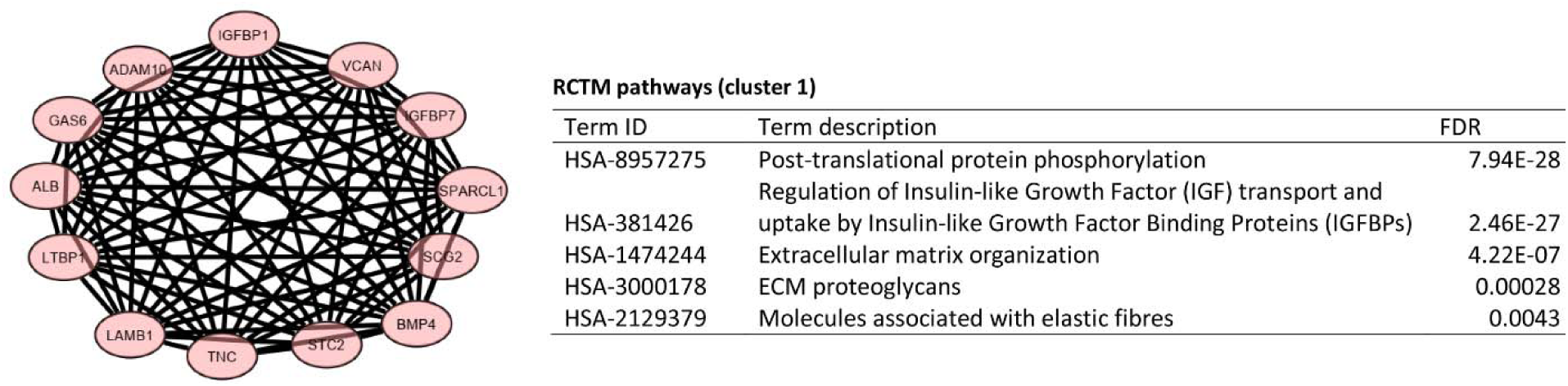
Top 1 module from the protein-protein interaction network between OA and RA.

**Figure 3.**
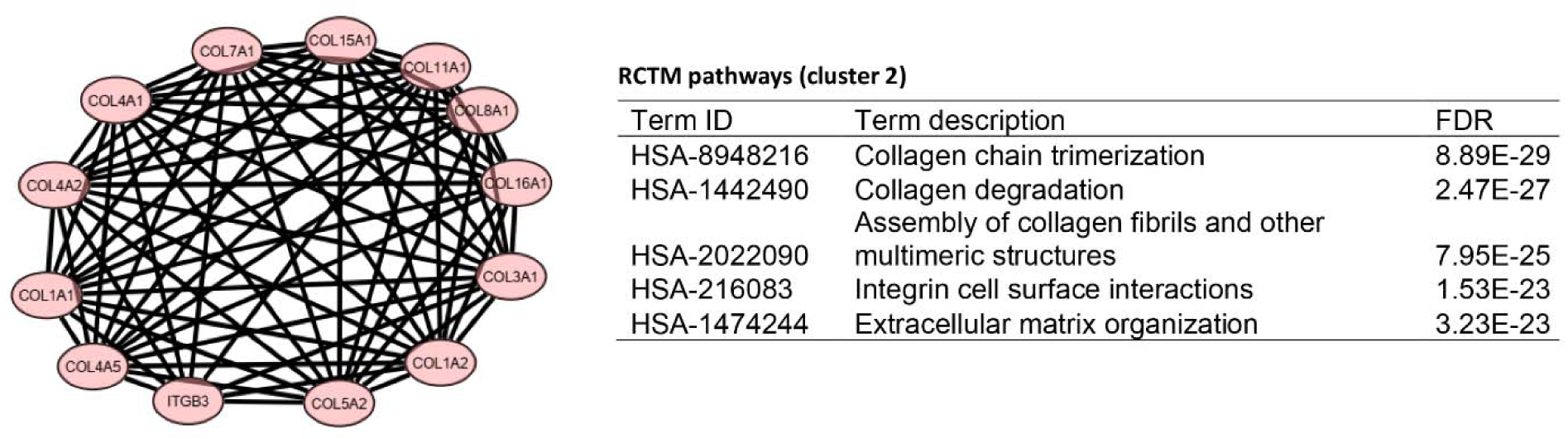
Top 2 module from the protein-protein interaction network between OA and RA.

## Discussion

The mechanisms of OA and RA show multiple overlaps, particularly in some enzymes and inflammatory regulators^16^. The chondrocytes and synovial cells bind to different cartilage components, which become a signaling scaffold regulating a number of bioactive factors to induce OA^22^. Then, the adjacent cells are activated to trigger cartilage degradation in OA and RA^23^. Thus, understanding of the different mechanisms of OA and RA will eventually provide therapies for both diseases.

Previous studies have shown that inadequate fluid, vascular disease, and ischemia can cause the osteocyte apoptosis, activation of osteoclasts^24, 25^. These areas of bone damages are commonly observed in OA with painful joints^26, 27^. Similar to our analysis, the KEGG analysis showed that the OA majorly affects the “Vascular smooth muscle contraction”, “Focal adhesion” and “ECM-receptor interaction” in comparison to RA. These suggest that the development of OA is more prone to angiogenesis or blood circulation.

In addition, OA is also associated with biological adhesion and cell adhesion in comparison to RA^28^. By GO analysis, we found most biological functions are relative to the adhesion processes including "Growth factor binding", "Extracellular matrix structural constituent" and "Integrin binding". Recently, pathophysiologic mechanism of OA has been put emphasis on the interactions between cartilage and synovium^5, 29^. Such concepts show a role not only of inflammatory markers in the pathogenesis of OA but also of adhesion molecules which regulates cell–cell interactions^30, 31^. It was proved by Georg Schett et al. that Vascular cell adhesion molecule 1 (VCAM‐1) is as a predictor of severe osteoarthritis of joints^32^. Thus, it is possible that the biological adhesion plays more important roles in OA than those in RA.

In our study, the PPI network identified several DEGs as potential critical genes which could be considered as drug targets or biomarkers^33^. Human Albumin has been shown to relieve pain and inflammation in OA^34^. Mitogen-Activated Protein Kinases (MAPK3) that involves different pathways is proved to be as a therapeutic target in OA^35^. Protein tyrosine phosphatase receptor type C (PTPRC) is a critical predictor of response to anti-TNF therapy in OA^36^. The mutations in COL1A1 gene can cause histology changes of subchondral bone that further lead to OA. CXCL12 can regulate aggrecanase activation and cartilage degradation in a post-traumatic OA^37^. Periostin is related to prevalent knee OA in women^38^. Collagen type III alpha 1 chain (COL3A1) can affect the ER stress in the vascular system^39, 40^. Loeser et al. reported significant elevation of COL3A1 in an OA mice model^41^. NCAM1 plays a crucial role in diverse functions in osteogenesis and chondrogenesis^42^. Biglycan was expressed in cartilage and bone, which can contribute to the accelerated OA in the temporomandibular joint^43^. Our analysis further indicated that macromolecule metabolism and collagen metabolism signaling pathways are involved in OA in comparison to RA by analyzing the Reactome Pathway Database.

In summary, our study provided the basis for the identification of difference biomarkers between OA and RA. Biological adhesion and cell adhesion are two key points affected by OA. Future studies will focus on the specific and precise OA drugs on clinical trials. This study thus provides further insights into the characteristics of OA, which may facilitate the accurate diagnosis and drug development.

